# Cerebellar Activation Bidirectionally Regulates Nucleus Accumbens Core and Medial Shell

**DOI:** 10.1101/2020.09.28.283952

**Authors:** Alexa F. D’Ambra, Ksenia Vlasov, Se Jung Jung, Swetha Ganesan, Evan G. Antzoulatos, Diasynou Fioravante

**Affiliations:** Center for Neuroscience, Davis CA 95616, United States; Department of Neurobiology, Physiology and Behavior, University of California Davis, Davis CA 95616, United States

**Author notes:** These authors share equal authorship (Ksenia Vlasov, Se Jung Jung). These authors share senior authorship (Evan G. Antzoulatos, Diasynou Fioravante). Gillings School of Global Public Health, University of North Carolina (Swetha Ganesan). Correspondence should be addressed to: Diasynou Fioravante, PhD.

**Keywords:** Cerebellum, nuclei, nucleus accumbens, ventral striatum, thalamus, intralaminar, VTA, connectivity, mouse, optophysiology, circuit mapping, optogenetics, motivated behavior

## Abstract

Although the cerebellum is now recognized as part of a long-range brain network that serves limbic functions and motivated behavior, knowledge of cerebello-limbic connectivity is limited, and nothing is known about how the cerebellum connects functionally to the nucleus accumbens (NAc). Here, we report that stimulation of cerebellar nuclei in mice of both sexes modulates spiking activity in both NAc core and medial shell with fast excitation and slower, less synchronized inhibition. Fast responses would be well poised to support rapid communication of information critical to the control of motivated behavior, whereas slower responses may be suggestive of a regulatory function, such as gain control. Tracing experiments to chart cerebellar nuclei-NAc pathways identified disynaptic pathways that recruit the ventral tegmental area (VTA) and intralaminar thalamus (Centromedial and Parafascicular nuclei) as intermediary nodes. Optogenetic activation of cerebellar axons in each of these nodes was sufficient to evoke responses in both NAc core and medial shell, albeit with distinct, node-dependent properties. These pathways and the functional connectivity they support could underlie the role of the cerebellum in motivated behavior.

## Introduction

The cerebellum (CB) exploits the modular organization of its circuitry to integrate information and perform complex computations. Most cerebellar research sheds light on this complexity in the context of updating and predicting motor movements (Ito, 2006). However, accumulating evidence supports cerebellar involvement in high-order non-motor functions as well (Buckner, 2013; Sokolov et al., 2017; Liang and Carlson, 2019; Hull, 2020). In humans, the CB is consistently activated during decision making, particularly during risk- or reward-based tasks (Ernst, 2002; Guo et al., 2013), and during aversive experiences and emotional learning (Moulton et al., 2014; Ernst et al., 2019). Further support for non-motor roles of the CB stems from clinical translational studies, which have linked CB dysfunction with neurodevelopmental disorders, posttraumatic stress disorder, generalized anxiety disorder, addiction, and cognitive and emotional disturbances known as cerebellar cognitive affective syndrome (Roy et al., 2013; Volkow et al., 2013; Moulton et al., 2014; Wang et al., 2014; D’Mello and Stoodley, 2015; Miquel et al., 2016; Lanius et al., 2017; Rabellino et al., 2018; Sathyanesan et al., 2019; Schmahmann, 2019). These findings are further corroborated by evidence from animal studies, which solidify a role for the CB in the processing of valence, aversion, reward, reward anticipation and omission (Wagner et al., 2017; Carta et al., 2019; Heffley and Hull, 2019; Kostadinov et al., 2019; Ma et al., 2020; Shuster et al., 2021); emotional learning and aggression (Supple et al., 1987; Sacchetti et al., 2002; Lorivel et al., 2014; Strata, 2015; Otsuka et al., 2016; Adamaszek et al., 2017; Frontera et al., 2020; Jackman et al., 2020); and motivation (Caston et al., 1998; Bauer et al., 2011; Peterson et al., 2012; Baek et al., 2022).

The ability of the CB to derive a complex repertoire of non-motor functions from its relatively invariant cellular organization is largely attributed to its diverse outputs (Reeber et al., 2013; Xiao and Scheiffele, 2018; Fujita et al., 2020; Judd et al., 2021). In addition to the heavily emphasized non-motor cerebellar influences on cortical regions (Ito, 2008; Mittleman et al., 2008; Strick et al., 2009; Watson et al., 2014; Gao et al., 2018; Brady et al., 2019; Kelly et al., 2020), functional and/or anatomical cerebellar connections with subcortical structures that are critical for cognition and emotion have also been documented (Snider and Maiti, 1976; Heath et al., 1978; Dietrichs and Haines, 1989; Rochefort et al., 2011; Fujita et al., 2020; Zeidler et al., 2020; Jung et al., 2022). Here, we focused on the nucleus accumbens (NAc), a key limbic structure that shares reward, motivation and affective functionality with the CB (Floresco, 2015; Klawonn and Malenka, 2018). The core and medial shell (NAc_Core_, NAc_Med_) subregions of NAc exhibit distinct input-output organization, anatomy, and function (Groenewegen et al., 1999; Zahm, 1999; Corbit and Balleine, 2011; Ito and Hayen, 2011; Badrinarayan et al., 2012; Li et al., 2018), with roles in social behavior and reward learning that are often opposing (Floresco et al., 2018; Shan et al., 2022). Thus, the investigation of CB connections to NAc subregions could offer insights into the complexities of CB limbic functionality. Stimulation of CB cortex or deep cerebellar nuclei (DCN) modulates levels of NAc dopamine -- an important, but not exclusive, regulator of NAc functions (Dempesy et al., 1983; Albert et al., 1985; Holtzman-Assif et al., 2010; Luo et al., 2018; Holloway et al., 2019; Root et al., 2020; Zell et al., 2020; Low et al., 2021). However, how the CB connects to NAc functionally is unknown.

Here we used in vivo electrophysiology in anesthetized mice to examine the effects of DCN stimulation on spiking activity in the NAc. We provide the first evidence, to our knowledge, of an electrophysiological connection between CB and NAc, which shows NAc subregion-dependent specificity. Using viral tracing approaches, we offer insights on an anatomical blueprint of disynaptic CB-NAc connectivity, which includes nodal neurons in VTA and limbic thalamus (Centromedial nucleus, CM; and Parafascicular nucleus, PF). Through optogenetic stimulation of DCN projections to each of these nodes, we uncover circuit-dependent connectivity features that are NAc subregion-specific. These findings offer foundational insights into the contribution of the CB to limbic functions such as motivation, reward learning, and affective processing.

## Results

### DCN microstimulation bidirectionally modulates NAc spiking activity

To examine functional connectivity between CB and NAc, we recorded ongoing spiking activity from 546 putative single units (N = 42 mice) in NAc *in vivo* while electrically microstimulating the DCN (Fig. 1A). We primarily targeted the lateral DCN, activation of which has been shown to modulate levels of dopamine in NAc (Holloway et al., 2019; Low et al., 2021). In a subset of experiments, post-hoc analysis localized the bipolar stimulating electrode in the interposed DCN, and these data were included in our analyses. Stimulation of DCN with five 100-μA current pulses at 200 Hz (total stimulus duration: 25 ms) evoked bidirectional modulation of NAc spiking activity (Fig. 1B). Across all recorded units, average spiking initially increased to 12.8 ± 0.4 spikes/s from a pre-stimulus baseline of 10.4 ± 0.5 spikes/s, followed by a modest, slow-recovering decrease to 9.8 ± 0.3 spikes/s. Because of jitter in response latency across neurons, the average spiking rate is an underrepresentation of the true magnitude of modulation, which is more accurately reflected in the average maximum activity. In response to stimulation, average maximum activity increased to 20 ± 0.6 spikes/s for all recorded single-units (t-test for paired samples: t_1017_ = -33; p < 0.001), followed by a decrease to an average minimum of 4 ± 0.3 spikes/s (t_1017_ = 35.7; p < 0.001) (n = 1018 units x stimulation sites). Further analysis identified three distinct response-types: excitatory (defined as firing rate increase >3σ above the baseline) (Fig. 1C), inhibitory (defined as firing rate decrease >1σ below the baseline for 3 consecutive time bins) (Fig. 1D), and mixed (both excitatory and inhibitory) (Fig. 1E).

**Figure 1.**
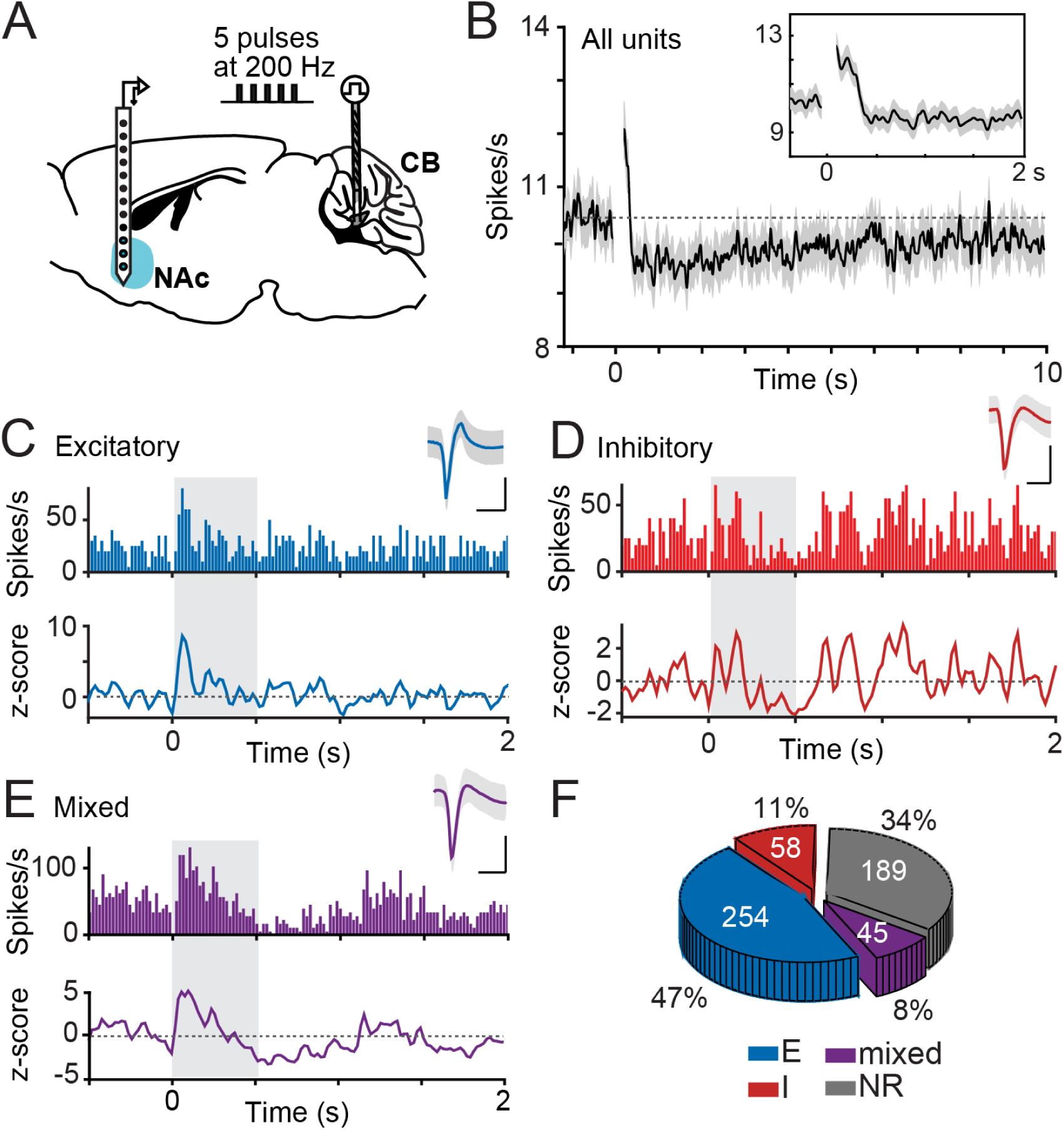
Cerebellar stimulation effectively elicits excitatory and inhibitory responses in nucleus accumbens. **A**, Schematic diagram of recording setup. Stimulation protocol consisted of 10 trials (inter-trial interval: 15 s) of five 100-µA, 0.5-ms pulses at 200 Hz. NAc: nucleus accumbens; CB: cerebellum. **B**, Average spiking activity (spikes/s) across all recorded units (N = 42 mice). Inset shows zoom-in of first two seconds after stimulation. **C**,**D**,**E**, Examples of excitatory (**C**), inhibitory (**D**), and mixed (**E**) modulation in the NAc. *Top,* Peri-stimulus time histogram (PSTH) (10-ms bins) of the average firing rate across 10 trials. *Bottom,* Baseline-normalized firing rate (z-score) across time. *Inset*, Example average waveform of putative single unit. Scale bars: 1 ms, 10 µV. Shaded area marks response window. **F,** Distribution of response types elicited in NAc by DCN stimulation. *Blue*: Excitatory (E), *Red*: inhibitory (I), *Purple:* mixed, *Gray*: non-responders (NR). Number of recorded units per response-type (*white*) and corresponding percentages are shown.

Of the 546 recorded NAc neurons, 66% responded to DCN stimulation, with most (47%) displaying an excitatory response (Fig. 1F). An inhibitory response was elicited in 11% of neurons, whereas a smaller fraction (8%) displayed mixed responses. The average excitatory response amplitude was 349 ± 15% of baseline, and the average inhibitory response amplitude was 30 ± 2% of baseline.

### Effects of DCN microstimulation on NAc core and medial shell spiking activity

The NAc is divided into two distinct subregions: a core subregion (NAc_Core_) surrounded by an outer subregion, the medial aspect of which is known as the medial shell (NAc_Med_). Each of these subregions has unique microanatomical, functional and connectivity features (Floresco, 2015; Salgado and Kaplitt, 2015). We aimed to examine whether the difference in proportions (i.e., prevalence) between DCN-evoked excitatory and inhibitory responses in NAc stemmed from regional differences (core vs. medial shell) at NAc recording sites. Our axial multi-electrode array approach enabled recordings from multiple NAc sites in both NAc_Core_ and NAc_Med_ (Fig. 2A). Out of 169 neurons we recorded in NAc_Core_, more than half (53%) responded with excitatory modulation to 100-µA stimulation of at least one site in DCN, while 8% responded with inhibitory modulation, and another 8% with mixed modulation (Fig. 2B *left*). Similarly, out of 377 neurons we recorded in NAc_Med_, 44% responded to DCN stimulation with excitation, 12% with inhibition, and 8% with mixed modulation of spiking activity. The frequency distributions of these response-types in NAc_Core_ and NAc_Med_ were not significantly different (χ^2^_(3)_ = 4.7, p = 0.19).

**Figure 2.**
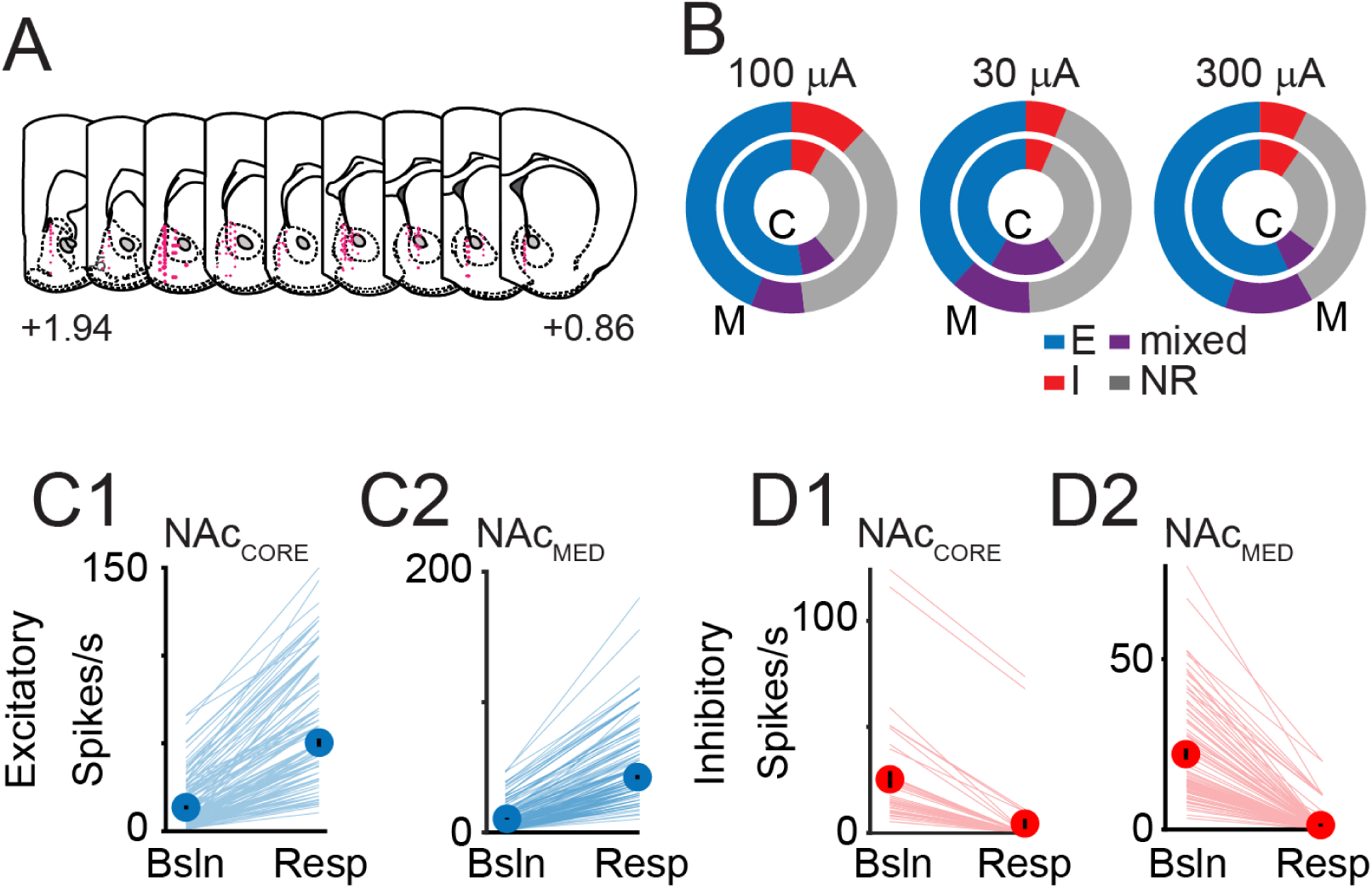
Nucleus accumbens responses to cerebellar stimulation. **A**, Recording sites in nucleus accumbens core (NAc_Core_) and medial shell (NAc_Med_). Numbers indicate distance (in mm) from bregma. **B,** Distribution of response types elicited by 30-, 100-, or 300-µA DCN stimulation. N_30_ = 14 mice, N_100_ = 42 mice, N_300_ = 40 mice. Outer ring (M): NAc_Med_; inner ring (C): NAc_Core_. *Blue:* excitatory (E), *Red*: inhibitory (I), *Purple:* mixed, *Gray:* non-responders (NR). **C,** Comparison between average baseline (Bsln) and peak firing rate (Resp) elicited by 100-µA DCN stimulation for excitatory responses in NAc_Core_ (**C1**) and MAc_Med_ (**C2**). Group averages (± SEM) are overlaid onto lines representing single units. **D1**,**2**, Same as C but for inhibitory responses.

We then examined whether the amplitude of excitatory and/or inhibitory responses varied between NAc_Core_ and NAc_Med_ (Figs. 2C-D). For this analysis, the excitatory and inhibitory components of mixed responses were considered together with the excitatory and inhibitory response types, respectively. In both NAc_Core_ and NAc_Med_, DCN stimulation elicited significant excitatory modulation of firing rate (mean ± sem, spikes/s in NAc_Core_: baseline: 12.2 ± 1.1, peak excitatory response: 52.2 ± 2.5, paired t_162_ = -24.2, p < 0.0001; in NAc_Med_: baseline: 7.9 ± 0.6, peak excitatory response: 42.2 ± 1.5, paired t_267_ = -32.4, p < 0.0001), which did not differ between subregions (% baseline: NAc_Core_: 856.2 ± 79.2, NAc_Med_: 978.8 ± 62.5, t_429_= -1.2, p = 0.23). Similarly, both NAc_Core_ and NAc_Med_ showed significant inhibitory modulation (mean ± sem, spikes/s in NAc_Core_: baseline: 24.3 ± 3.9, peak inhibitory response: 4.0 ± 2.3, paired t_42_ = 9.7, p < .0001; in NAc_Med_: baseline: 22.1 ± 1.6, peak inhibitory response: 1.3 ± 0.4, paired t_92_ = 15.5, p < 0.0001), which did not differ between subregions (% baseline: NAc_Core_: 4.3 ± 2.1, NAc_Med_: 2.6 ± 0.8, t_134_ = 0.9, p = 0.36)(p-values corrected for multiple comparisons). These results suggest that cerebellar signals are equally likely to reach NAc_Core_ and NAc_Med_. Upon further inspection, we found that irrespective of subregion, the baseline firing rate of excitatory responses was significantly lower than the baseline of inhibitory responses (F(1,563) = 88, p < 0.001). Differences in baseline between excitatory and inhibitory responses could arise from potentially different NAc cellular properties, although the fact that neurons with elevated firing rates are more likely to satisfy our criterion for inhibitory responses should also be considered.

### Differential and non-linear dependence of NAc responses on DCN stimulation intensity

To further probe the functional connectivity between DCN and NAc, we assessed the effectiveness of DCN microstimulation at two more intensities, 30 μA and 300 μA, in addition to 100 μA (Fig. 2B; *middle* and *right*). In NAc_Core,_ the proportion of excitatory responses changed from 42% (with 30 μA stimulation) to 53% (with 100 μA), to 57% (with 300 μA); the proportion of inhibitory responses changed from 7% (30 μA) to 8% (100 μA) to 10% (300 μA); and finally, the proportion of mixed responses changed from 18% (30 μA) to 8% (100 μA) to 8% (300 μA). Contingency table analysis did not find a significant effect of stimulation intensity on the frequency distribution of response-types in NAc_Core_ (χ^2^_(6)_ = 9.4; p = 0.15). In NAc_Med_, the proportion of excitatory responses changed from 38% (30 μA) to 44% (100 μA) to 45% (300 μA); the proportion of inhibitory responses changed from 6% (30 μA) to 12% (100 μA) to 7% (300 μA); and the proportion of mixed responses changed from 13% to 8% to 14%. The effect of stimulation intensity on the frequency distribution of response-types in NAc_Med_ was statistically significant (χ^2^_(6)_ = 13.6; p < 0.05), suggesting that the NAc subregions displayed differential sensitivity to the intensity of DCN microstimulation. For both NAc subregions, the prevalence of the different response-types did not change proportionally to the ∼3-fold change in stimulation intensity, which suggests non-linear dependence to stimulation intensity.

### The bidirectional NAc modulation is not a reflection of topographical DCN organization

In our experiments we used bipolar stimulation electrodes to contain the spread of electrical current at DCN stimulation sites. To confirm that the stimulation was indeed localized, we examined the likelihood of NAc neurons that responded to stimulation of individual DCN sites to also respond to adjacent stimulation sites along anatomical axes in the same experiment (Fig. 3A1-2). We found that the probability of a NAc_Core_ neuron that responded to stimulation of a DCN site to also respond to stimulation of a site ∼100 μm away dropped by 66%. In NAc_Med_, this probability dropped by 33%. The subregion-dependent difference in the likelihood of adjacent DCN sites to evoke a response in the same neuron most likely reflects differences in the architecture of the neural circuits. Regardless of the difference between NAc subregions, the probability decrease is consistent with localized stimulation.

Given that DCN stimulation appeared spatially restricted, could the observed differences in NAc response-types arise from regional differences within DCN? To address this question, we examined whether there was topographical clustering of DCN sites with respect to their effectiveness to evoke significant excitatory or inhibitory responses in NAc_Core_ and/or NAc_Med_. We did not find indications of such clustering (Fig. 3B1-3, C1-3), which argues against topographical organization of lateral/interposed DCN with respect to NAc response types. Using a similar approach, we examined whether there was topographical clustering of sites with excitatory and/or inhibitory responses within NAc subregions, but we did not find evidence for any such organization either (not shown).

**Figure 3.**
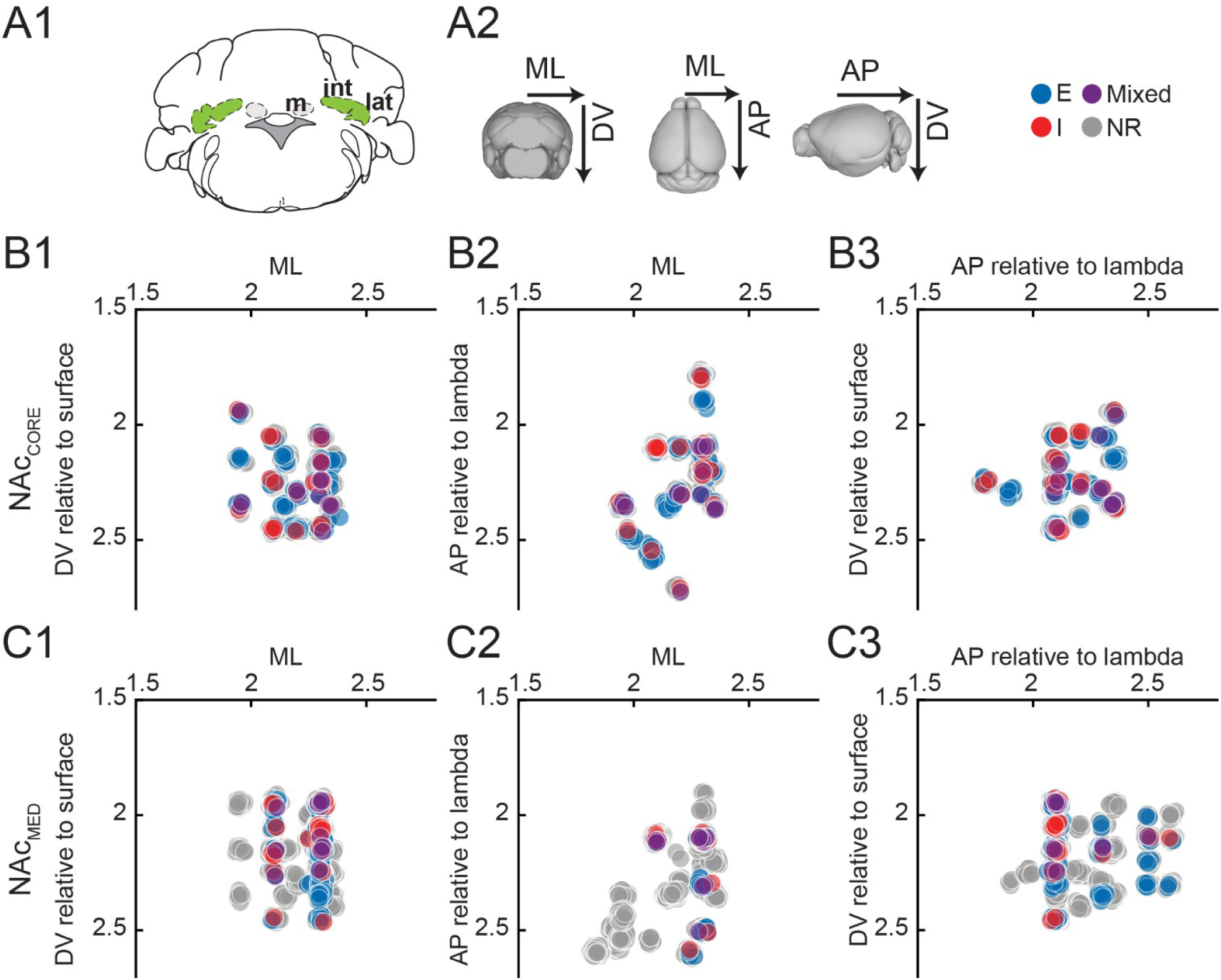
The distribution of response-types in nucleus accumbens subregions is not due to topographical specialization within deep cerebellar nuclei. A, The lateral (lat) or interposed (int) DCN (A1) was electrically stimulated. M: medial n. A2, Demarcation of anatomical planes: ML: medio-lateral; DV: dorsoventral; AP: antero-posterior. B,C, Stereotactic coordinates of DCN stimulation sites in ML-DV (B1,C1), ML-AP (B2,C2), and AP-DV (B3,C3) planes. Colored dots denote sites that evoked excitatory (excit.: *blue*), inhibitory (inh.: *red*), mixed (*purple*) or no responses (NR: *gray*) in NAc_Core_ (B1-3) and NAc_Med_ (C1-3).

### Anatomical blueprint of DCN-NAc connectivity

There are no direct, monosynaptic connections between DCN and NAc (Allen Brain Atlas, and our own observations); we therefore hypothesized the existence of at least disynaptic anatomical pathways between the two areas. To test this hypothesis, we adopted a 2-pronged approach. First, we combined injection of a retrograde tracer (cholera toxin subunit B (ctb) -640 or -568) in NAc with injection of an anterograde viral tracer (AAV9-CAG-GFP) in DCN to identify areas of overlap (nodes) in a putative disynaptic CB-NAc circuit (Fig. 4A,B) (N = 6 mice). Histological processing and high-resolution confocal imaging of brain sections revealed two regions of overlap between ctb-filled neurons that project to NAc and GFP- labelled DCN axonal projections: the VTA and limbic thalamus (Fig. 4C,D). In VTA, most areas of overlap localized medially and caudally and involved both TH+ (dopaminergic) neurons (Fig. 4C) and TH- neurons. In the thalamus, we found areas of overlap in centromedial (CM) and parafascicular (PF) intralaminar nuclei (Fig. 4D). We found overlap in these same regions also when DCN injections were restricted to the lateral cerebellar nucleus only (Fig. 4E,F). These results are consistent with the existence of a disynaptic DCN-NAc anatomical circuit and point to VTA, and CM and PF thalamic nuclei, as putative circuit nodes.

**Figure 4.**
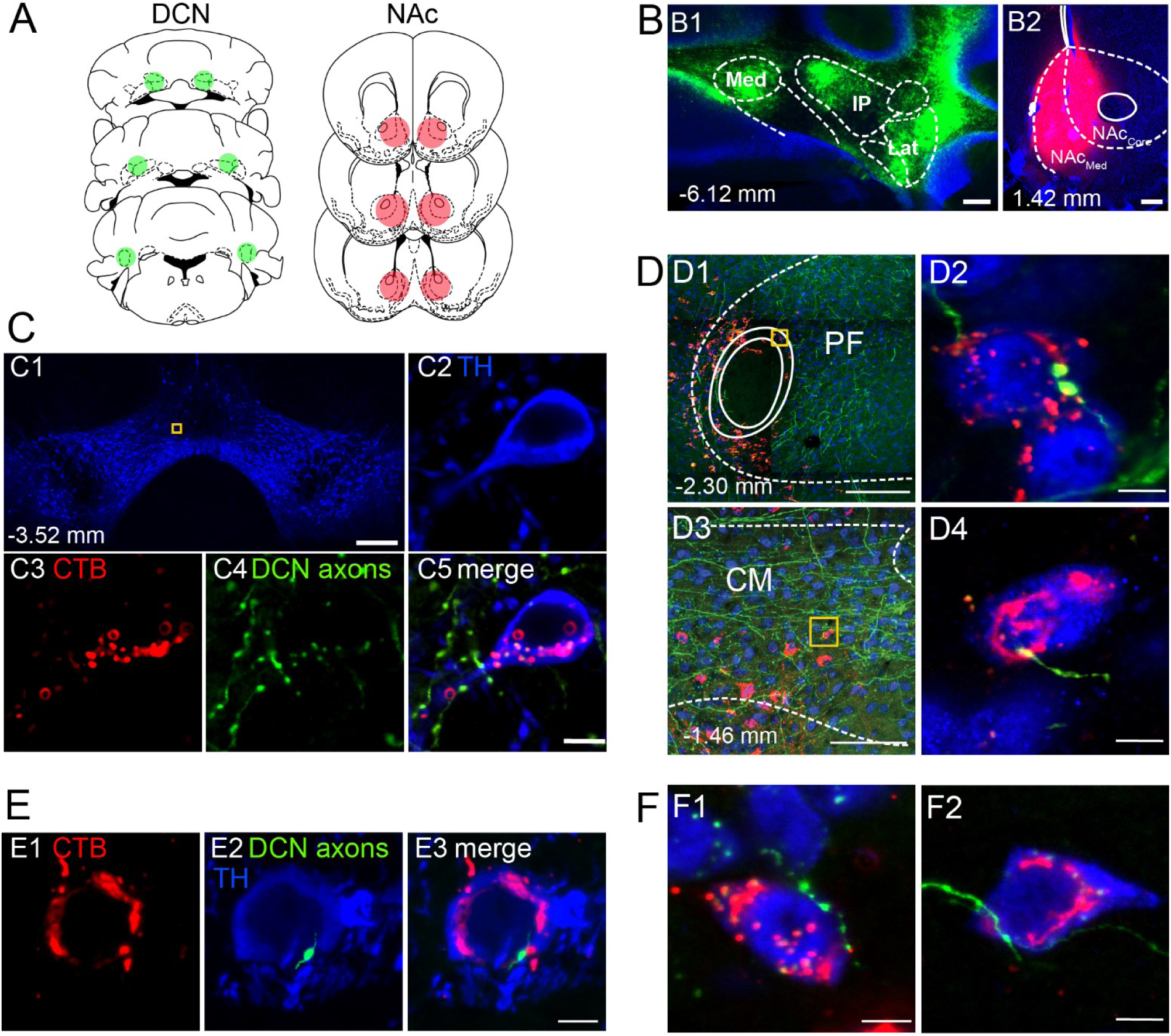
Co-localization of nucleus accumbens-projecting neurons with cerebellar projections in VTA and intralaminar thalamus. **A**, Schematic diagrams of stereotactic injections in DCN and NAc. **B1**, Expression of GFP at DCN injection sites. **B2**, Ctb-Alexa 568 injection site in NAc. C, Overlap of ctb-labeled NAc projectors and DCN axons in VTA. **C1**, VTA identification through TH immunostaining. **C2**, TH+ neuron. **C3**, Retrograde ctb-Alexa 568 labelling in NAc-projecting cell. **C4**, GFP-expressing DCN axons. C5, C2-C4 merged. **D**, Overlap of ctb retrograde label (red) in NAc projectors (blue; NeuN) and GFP-expressing DCN axons (green) in intralaminar thalamic nuclei. **D1-D2**, Overlap in parafascicular (PF) n. **D3-D4**, Overlap in centromedial (CM) n. Yellow boxes in C1,D1,D3 denote zoom-in areas depicted in C2-C5, D2 and D4, respectively. **E-F**, Airyscan confocal images from experiments similar to **A** but with only lateral DCN injected with AAV-GFP. **E1**, Ctb-labeled (red) NAc-projecting cell in the VTA. **E2**, TH+ cell (blue) with overlapping GFP-labeled axonal projection (green) from lateral DCN. **E3**, E1-2 merged. **F1**, GFP-labeled axon from lateral DCN (green) overlaps with ctb-labeled (red) NAc-projecting cell (blue: NeuN) in the PF n. **F2**, Same as in F1 but for neuron in the CM n. Scale bars: B1-2,C1,D1: 200 µm; C2-C5,D2,D4,E1-3,F1,F2: 5 µm; D3: 100 µm. N = 6 mice. Numbers denote distance from bregma.

To confirm these observations through an independent approach, we performed AAV1- mediated anterograde transsynaptic tracing experiments (Zingg et al., 2017) via stereotactic injections of AAV1-Cre in DCN and AAV-FLEX-tdTomato in VTA or limbic thalamus (Fig. 5A,B) (N = 4 mice). This approach relies on the transsynaptic transfer of Cre to postsynaptic neurons, which, once infected with a floxed fluorophore, become fluorescently labeled. With this method, we confirmed the existence of neurons receiving CB input in VTA (Fig. 5C1,C2) and CM, PF intralaminar thalamus (Fig. 5D1,D2). Importantly, we were able to localize their labeled axonal projections in NAc, in both NAc_Core_ and NAc_Med_ (Fig. 5C3,D3). We observed similar patterns of cell labeling and projections to the NAc_Core_ and NAc_Med_ when AAV1-Cre was injected in lateral DCN only (Fig. 5E,F) (N = 3 mice). These findings chart a blueprint of disynaptic DCN-NAc connectivity and could provide an anatomical foundation for our newly discovered DCN-NAc functional connection.

**Figure 5.**
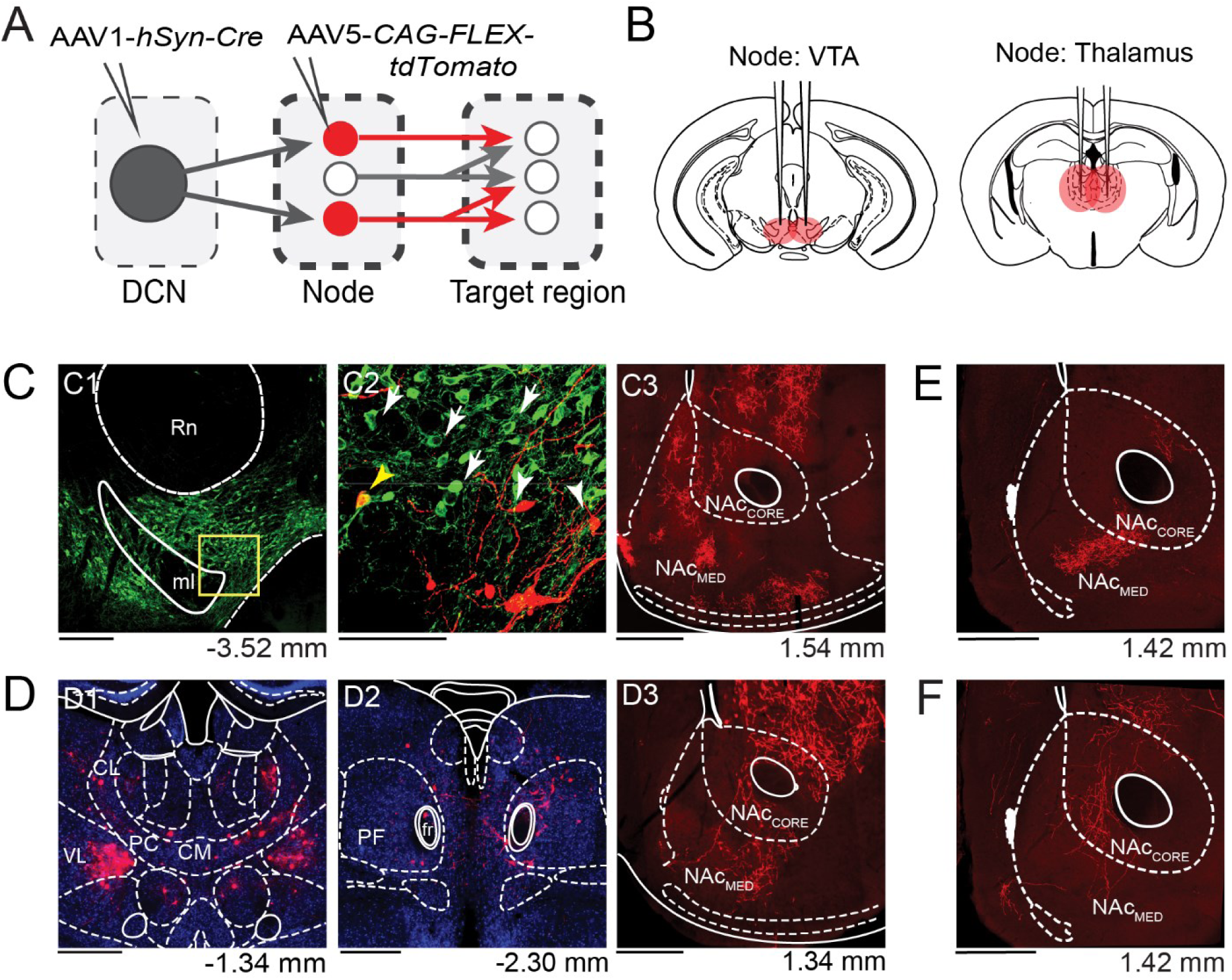
Disynaptic cerebellum - nucleus accumbens connectivity is confirmed via anterograde transsynaptic viral tracing. **A**, Schematic diagram of AAV1-mediated transsynaptic labeling approach. **B**, Schematic diagrams of stereotactic injections of floxed fluorophore in VTA and thalamic nodes. **C1**, The VTA is visualized via TH immunostaining (green). Yellow box denotes zoom-in area depicted in C2. Rn: red nucleus; ml: medial lemniscus. **C2**, Neurons that receive DCN input are labeled with tdTomato (red; white arrowheads). Green: TH+ neurons (white arrows). Yellow arrowhead: TH+, tdTomato+ neuron. **C3**, Projections of CB-VTA neurons in NAc_Core_ and NAc_Med_. **D1**, Neurons that receive DCN input in thalamus are labeled with tdTomato (red). Blue: NeuN. CM: centromedial n.; PC: paracentral n.; CL: centrolateral n.; VL: ventrolateral n. **D2**, Same as D1, but for parafascicular n. (PF). fr: fasciculus retroflexus. **D3**, Projections of CB-thalamic neurons in NAc_Core_ and NAc_Med_. **E**-**F**, Fluorescence images from experiments similar to A-B but with only lateral DCN injected with AAV1-Cre. **E**, NAc projections of VTA neurons receiving input from lateral DCN. **F**, Same as in E but for thalamic neurons. Scale bars below images: C1: 200 µm; C2: 100 µm; C3,D1- 3,E,F: 500 µm. N = 7 mice. Numbers denote distance from bregma.

### Photostimulation of DCN axonal projections in nodes supports the existence of disynaptic circuits between cerebellum and NAc

Our combined electrophysiological and neuroanatomical data point to the existence of neural circuits connecting the CB to the NAc_Core_ and NAc_Med_, with putative nodal regions in VTA and/or intralaminar thalamus. We sought to determine whether these anatomically defined nodes are also functional nodes serving CB-NAc connectivity. To this end, we performed additional electrophysiological experiments combined with optogenetic stimulation of DCN projections in the putative nodes (Figs. 6,7). We expressed channelrhodopsin (ChR2) in DCN projection neurons through viral injection (Figs. 6A,6B) and recorded responses in NAc_Core_ and NAc_Med_ following photostimulation of DCN axons (10 pulses at 20 Hz, 10 mW at fiber tip) in VTA (Fig. 6C1,C2), CM (Fig. 6D1,D2), or PF (Fig. 6E1,E2). All DCN nuclei were injected in order to maximize ChR2 expression and the chances of successful stimulation of projections.

**Figure 6.**
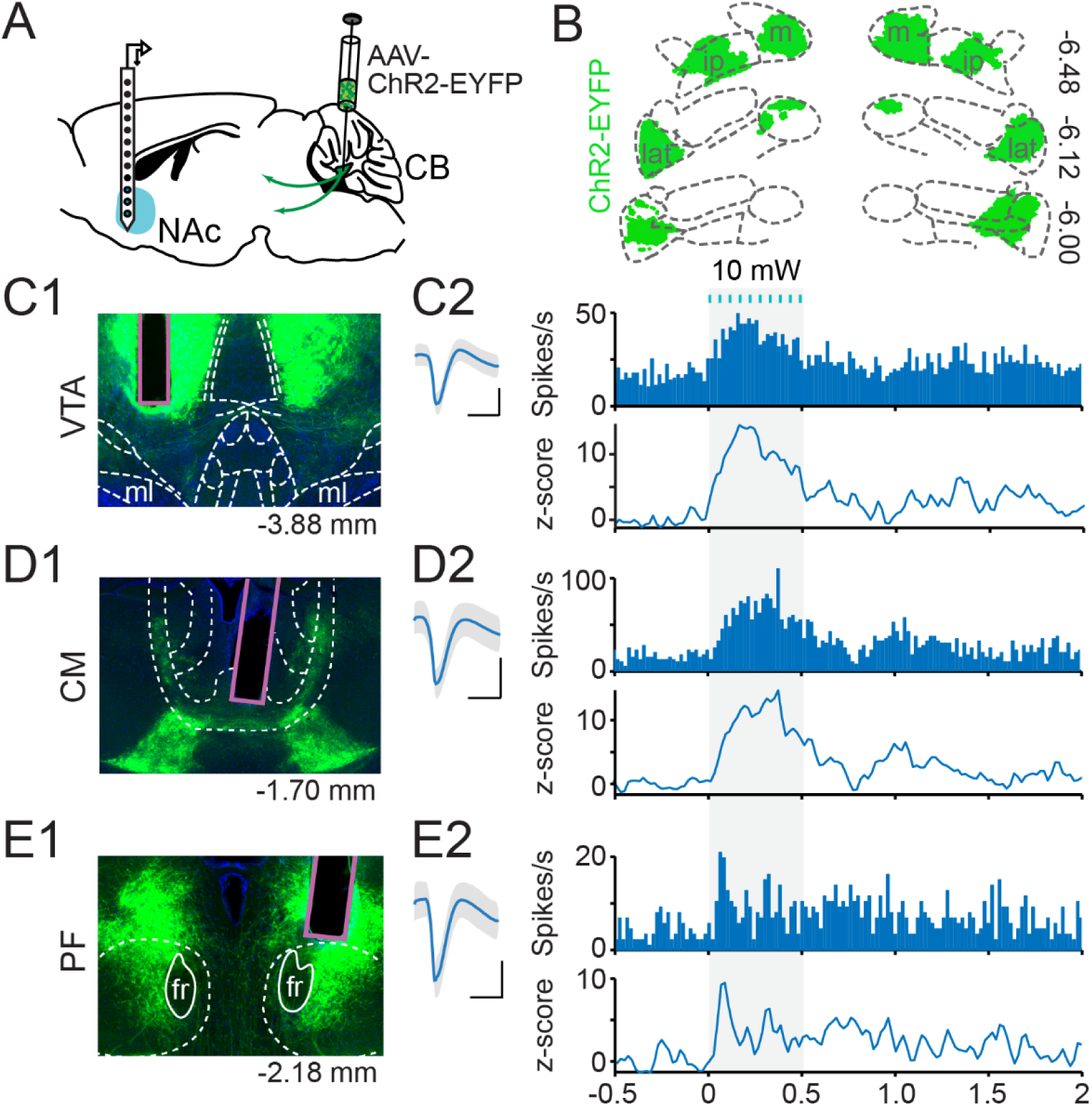
Photostimulation of cerebellar projections in VTA and thalamus elicits responses in nucleus accumbens. **A**, Schematic diagram of optophysiological approach. *NAc:* nucleus accumbens, *CB:* cerebellum. **B,** Example ChR2-EYFP expression in DCN. lat: lateral n.; ip: interposed n; m: medial n. **C**,**D**,**E**, Histology and NAc responses to optogenetic stimulation of VTA (**C1,2**), CM (**D1**,**2**), and PF (**E1**,**2**). **C1**,**D1**,**E1**: Representative images of optic fiber placement. ml: medial lemniscus; fr: fasciculus retroflexus. **C2**,**D2**,**E2**: Example single units. *Left*: Average spike waveform. Scale bars: 1 ms, 10 µV. *Top*: PSTH (10-ms bins) of firing rate (spikes/s). B*ottom*: Baseline-normalized firing rate (z-score) across time. Blue lines denote photostimulation; Shaded area marks response window. Numbers below or next to images indicate distance from bregma (in mm).

**Figure 7.**
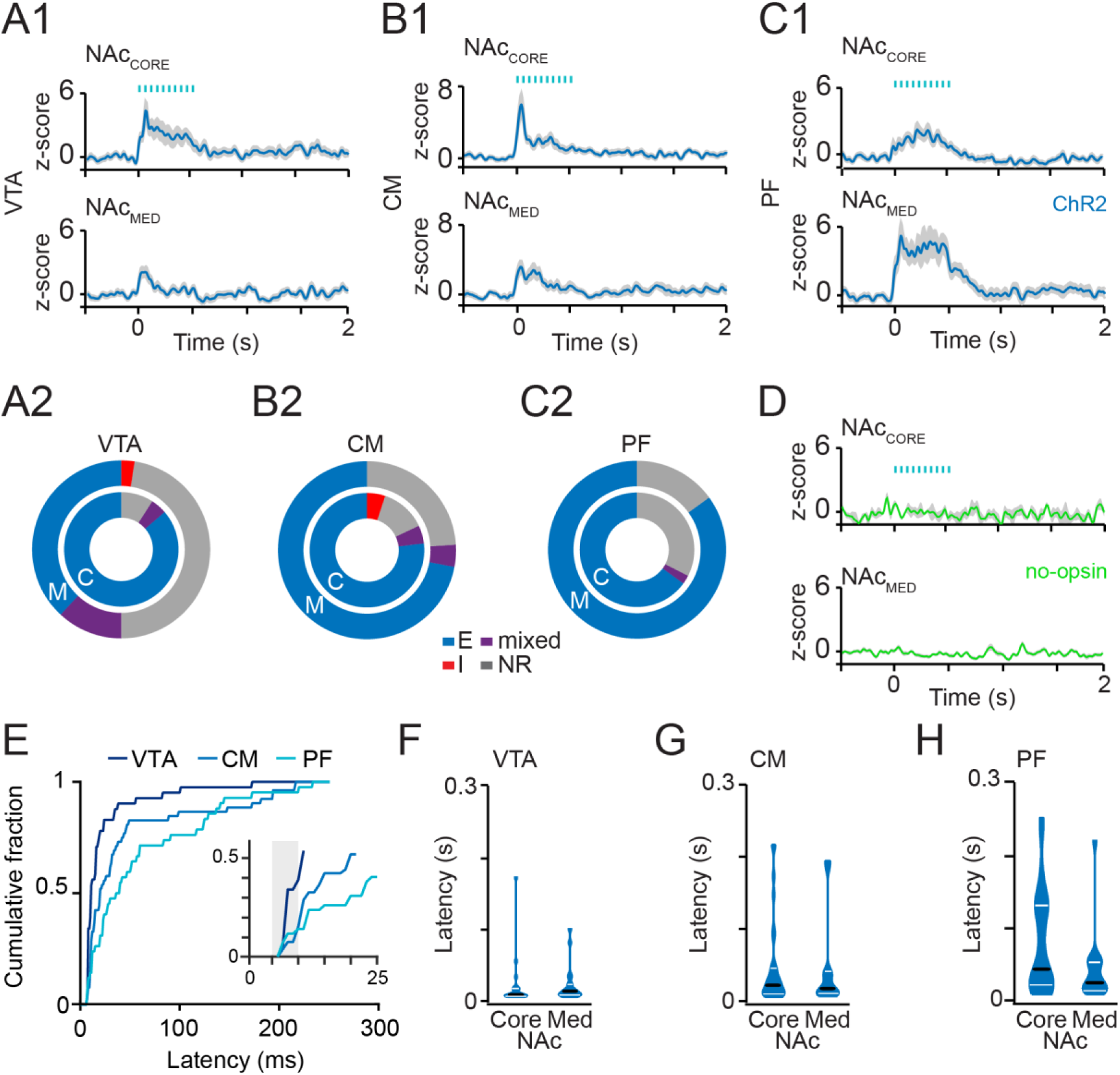
Node-dependent differences in nucleus accumbens responses to photostimulation of cerebellar projections. **A1**,**B1**,**C1**, Average normalized activity (z-score) in NAc_Core_ (*top*) and NAc_Med_ (*bottom*) as a function of time, for cerebellar fiber photostimulation (blue lines) in VTA (**A1**), CM (**B1**) and PF (**C1**). **A2**,**B2**,**C2**, Distribution of response-types in NAc_Core_ (C: inner ring) and NAc_Med_ (M: outer ring) upon photostimulation of cerebellar projections in VTA (**A2**), CM (**B2**), and PF (**C2**). *Blue:* excitatory (E), *Red*: inhibitory (I), *Purple:* mixed, *Gray:* non-responders (NR). **D**, Average normalized activity (z-score) in NAc_Core_ (*top*) and NAc_Med_ (*bottom*) as a function of time, for no-opsin controls. **E**, Cumulative histogram of NAc response onset latencies upon cerebellar fiber photostimulation in VTA, CM and PF. Inset: 0-25 ms zoom-in. **F**,**G**,**H**, Violin plots of onset latencies of excitatory responses in NAc_Core_ (Core) and NAc_Med_ (Med), upon cerebellar fiber photostimulation in VTA (**F**), CM (**G**) and PF (**H**). Black lines: Median; White lines: interquartile range (Q1-Q3). VTA: N = 4 mice, n_NAcCORE_ = 22 units, n_NAcMED_ = 42 units; CM: N = 4 mice, n_NAcCORE_ = 56 units, n_NAcMED_ = 26 units; PF: N = 3 mice, n_NAcCORE_ = 40 units, n_NAcMED_ = 20 units. D, no-opsin controls: N = 4 mice, n_NAcCORE_ = 5 units, n_NAcMED_ = 36 units.

As with electrical microstimulation of DCN sites, targeted photostimulation of ChR2- expressing DCN axons in each of the three putative nodes modulated spiking activity in both NAc_Core_ and NAc_Med_ (example units in Fig. 6C2,D2,E2; group averages in Fig. 7A1,B1,C1), establishing the VTA, CM and PF as functional nodes of the CB-NAc circuitry. Figure 7 shows the normalized group average excitatory responses of NAc_Core_ and NAc_Med_ neurons elicited by photostimulation in VTA (Fig. 7A1), CM (Fig. 7B1), and PF (Fig. 7C1). Because of jitter, the group average response underestimates the true magnitude of modulation, as discussed previously for electrical responses. Therefore, to further evaluate the connectivity strength in the three CB-NAc circuits, we computed the average peak amplitude of light-evoked excitatory responses in NAc subregions and asked whether it varied across nodes or between NAc subregions. In NAc_Core_, neurons displayed an average peak excitatory response to photostimulation in VTA of 43 ± 8 spikes/s; in CM: 44 ± 5 spikes/s; in PF: 36 ± 4 spikes/s. In NAc_Med_, the average peak excitatory response to photostimulation in VTA was 39 ± 6 spikes/s; in CM: 34 ± 3 spikes/s; and in PF: 41 ± 6 spikes/s. There was no statistically significant effect of NAc subregion, or of node, on peak responses (NAc subregion: F(1,120) = 0.4; p = 0.5; node: F(2,120) = 0.1; p = 0.9), and no significant interaction (F(2,120) = 0.9; p = 0.4). These NAc responses were specific to ChR2 expression, as no-opsin (EGFP-alone) controls did not show an effect of photostimulation on peak firing rate, even at 15 mW (response window vs. baseline: NAc_Core_: t_4_ = 0.53, p = 0.62; NAc_Med_: t_35_ = -1.41, p = 0.17) (Fig. 7D).

Light-evoked responses were excitatory in their majority, with frequency distribution that varied between NAc_Core_ and NAc_Med_ in a node-dependent manner: photostimulation of DCN projections in VTA evoked more excitatory responses in NAc_Core_ than in NAc_Med_ (excitatory responses in NAc_Core_: 86%, in NAc_Med_: 38%; χ^2^_(1)_ = 13.6; p < 0.001) (Fig. 7A2), whereas no such differences were observed upon photostimulation in CM (NAc_Core_: 77%, NAc_Med_, 72%; χ^2^_(1)_ = 0.20; p = 0.44) (Fig. 7B2) or PF (excitatory responses in NAc_Core_: 65%, in NAc_Med_ : 85%; χ^2^_(1)_ = 2.63; p = 0.11) (Fig. 7C2) (p-values corrected for multiple comparisons). Excitatory responses were also readily elicited in NAc_Core_ and NAc_Med_ with as little as 1 or 5 mW photostimulation (Table 1). Even at the lowest intensity (1 mW), a single pulse was sufficient to elicit a response (percent responses elicited by first pulse in VTA: NAc_Core_: 54%, NAc_Med_, 58%; in CM: NAc_Core_: 72%, NAc_Med_, 57%; in PF: NAc_Core_: 67%, NAc_Med_: 58%) (Table 2), pointing to high-fidelity connectivity that does not strictly require activity-dependent synaptic enhancement to propagate information to NAc. The proportion of responses elicited by the first light pulse tended to increase with photostimulation intensity more than expected by chance alone in VTA (permutation test: p < 0.001, n = 41, N = 4) and CM (p < 0.01, n = 52, N = 4), but not in PF (p = 0.08, n = 45, N = 3) (p-values corrected for multiple comparisons). This finding could potentially suggest convergent connectivity within the CB-VTA-NAc and CB-CM-NAc circuits.

**Table 1.**
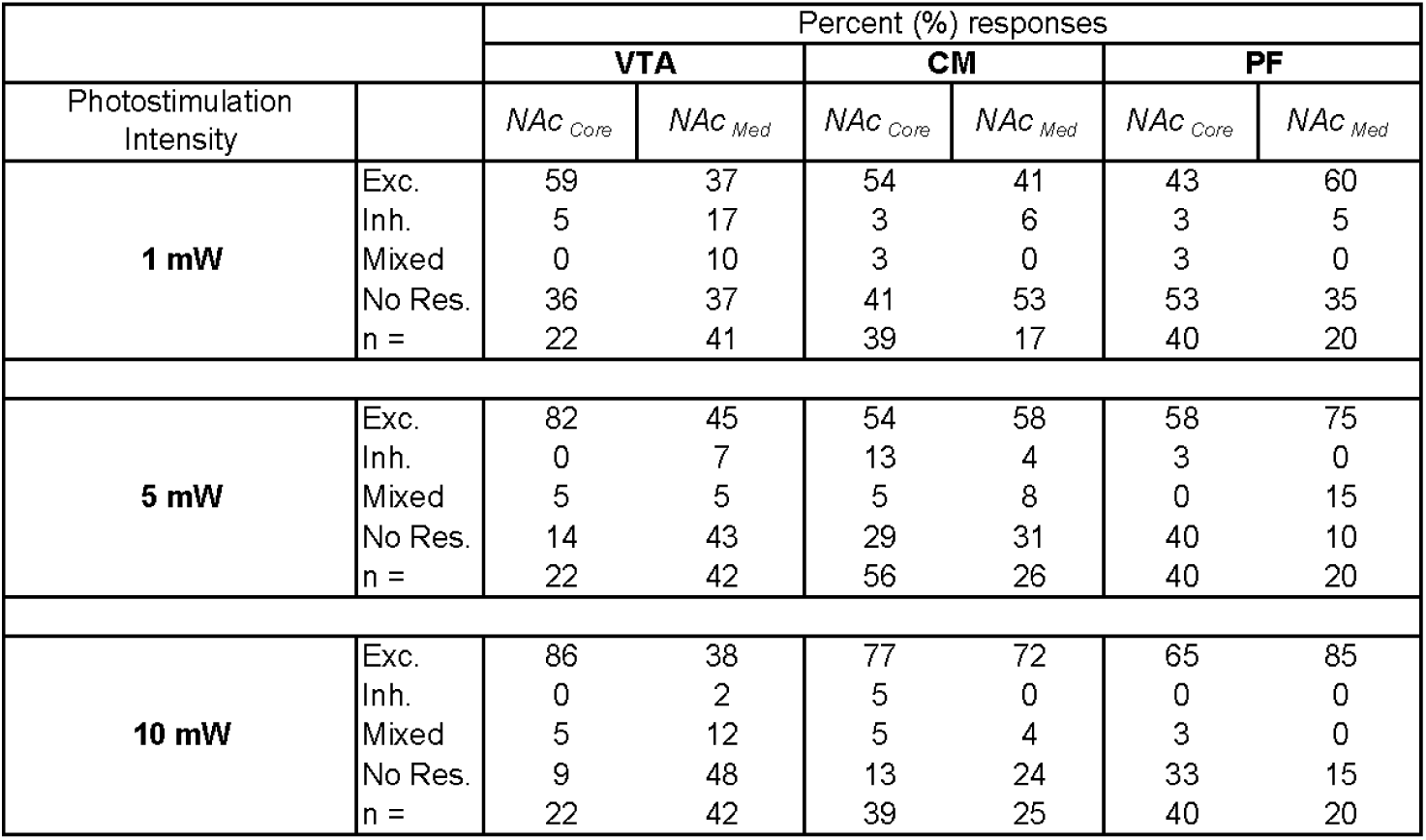
Percent responses in NAc_Core_ and NAc_Med_ elicited by photostimulation of cerebellar axons in VTA, CM and PF at different intensities.

**Table 2.**
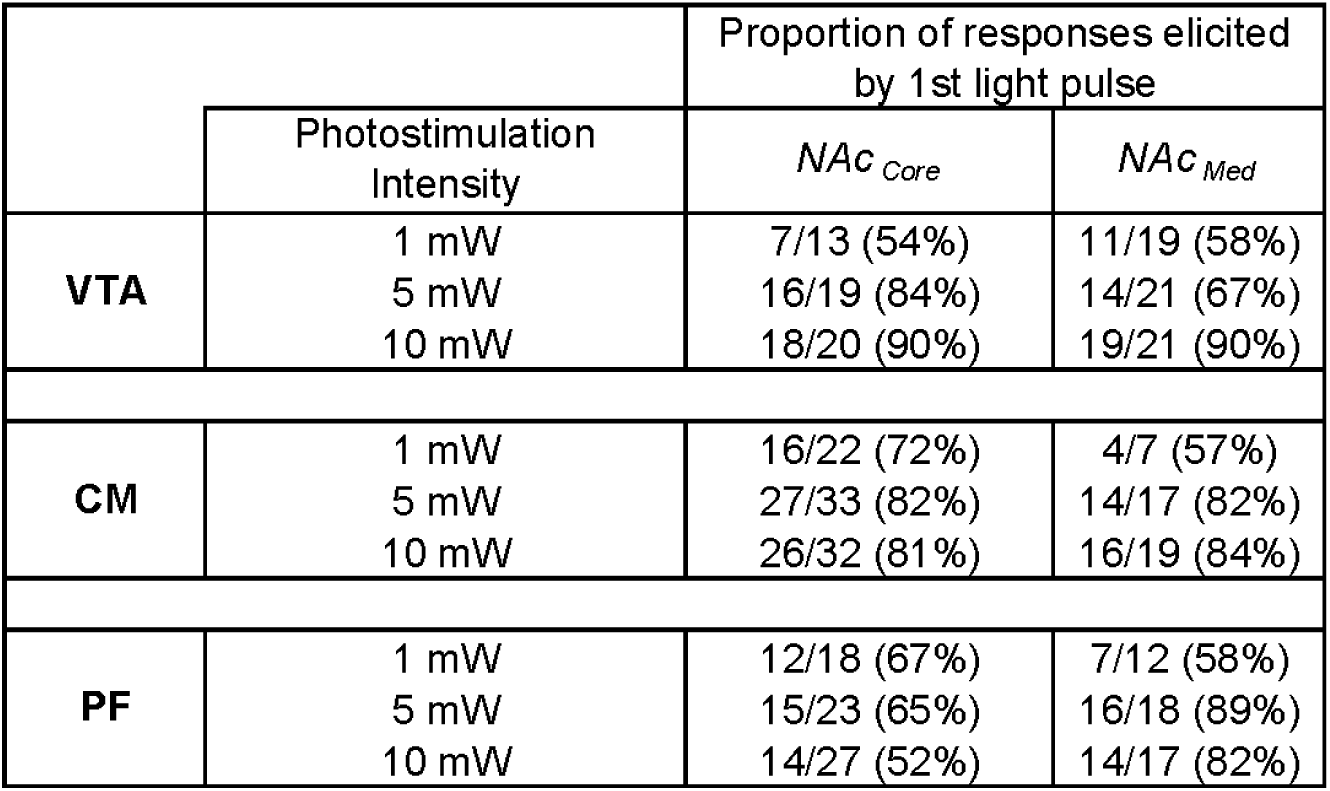
Proportion of excitatory responses in NAc_Core_ and NAc_Med_ elicited by the first light pulse, at different photostimulation intensities.

In contrast to excitatory responses, inhibitory responses were scarce in photostimulation experiments (Fig. 7A2, B2, C2). This finding appeared to be stimulation regime-dependent, because lowering the photostimulation intensity increased the prevalence of inhibitory responses in NAc more than expected by chance alone (permutation test, 1 vs. 5 vs. 10 mW: VTA: p < 0.01, n = 64, N = 4; CM: p < 0.05, n = 82, N = 4; PF: p < 0.05, n = 60, N = 3; p-values corrected for multiple comparisons) (Table 1). Finally, we asked whether each NAc subregion was differentially responsive to photostimulation in each of the nodes. We found no differential effect of 10 mW-photostimulation among the three nodes on the distribution of response-types in either NAc_Core_ (χ^2^_(2)_ = 5.4; p = 0.07) or NAc_Med_ (χ^2^_(2)_ = 4.9; p = 0.09).

### Temporal aspects of light-evoked responses

We wondered whether temporal aspects of CB-NAc communication differed between excitatory and inhibitory responses. This question cannot be addressed accurately in electrical stimulation experiments because of electrical artifacts, which can place a lower bound on the onset latency of detected responses, but can be readily addressed in optogenetic experiments. Analysis of onset latency of responses elicited by 10 mW CB axonal photostimulation indicated that, overall, excitatory responses had significantly shorter onset latencies than inhibitory ones (mean ± sem: excit.: 45.4 ± 5.0 ms, inh.: 256.7 ± 46.2 ms, t_147_= -10.0, p < 0.001; N = 10, n_exc._= 136, n_inh._= 13). To dissect node-specific effects, we computed the cumulative distributions of onset latencies for excitatory responses elicited by photostimulation in VTA, CM and PF (Fig. 7E). A relatively broad distribution of onset latencies was evident in all nodes, with half-rise latencies of 11 ms for VTA, 20 ms for CM and 40 ms for PF. Further analysis indicated that the probability of eliciting short-latency responses (≤ 10 ms), which are consistent with disynaptic connectivity (Chen et al., 2014; Gao et al., 2018; Assous et al., 2019; Yang et al., 2021), was greater for photostimulation in VTA compared to CM and/or PF (proportion of latencies ≤ 10 ms: VTA: 0.34, CM: 0.08, PF: 0.11; VTA vs CM: χ^2^_(1)_ = 10.3, p < 0.01; VTA vs PF: χ^2^_(1)_ = 6.4, p < 0.01; CM vs PF: χ^2^_(1)_ = 0.4, p = 0.54; p-values corrected for multiple comparisons) (Fig. 7E *inset*). Similar analysis could not be applied to inhibitory responses because of their limited prevalence (Fig. 7A2-C2 and Table 1). Finally, we examined NAc subregion-specific effects on excitatory response latencies (Fig. 7F-H). A two-way ANOVA corroborated the significant main effect of nodes (F(2,130) = 4.7; p < 0.05) but did not find significant main effect of NAc subregion (F(1,130) = 1.7; p = 0.19), or significant interaction effect between node and NAc subregion (F(2,130) = 1.7; p = 0.18).

Collectively, these results support and expand the conclusions of our neuroanatomical and electrical microstimulation studies that signals originating in DCN can reach NAc through pathways that involve the VTA and the intralaminar thalamus.

## Discussion

In this study we examined the previously uncharted functional connectivity between the cerebellum and nucleus accumbens using in vivo electrophysiology, neuroanatomy, and optogenetics. We found that electrical microstimulation of DCN elicits responses in both NAc_Core_ and NAc_Med_, manifested as excitatory and/or inhibitory modulations of neural firing rate. Neuroanatomical studies identified three disynaptic pathways through which cerebellar signals can reach the NAc. These pathways go through the VTA or the intralaminar thalamus (centromedial or parafascicular nucleus). Optogenetic activation of DCN axons in each of these intermediary nodes revealed that all three pathways can communicate cerebellar signals to NAc, albeit with distinct node-dependent properties.

### Bidirectional regulation of NAc through fast excitation and slow inhibition

Results from both electrical microstimulation and optogenetic experiments converged on the conclusion that the cerebellum sends strong excitatory signals to NAc. The strength of these excitatory signals did not show any NAc subregion-, or node-, dependence. However, the relative frequency of NAc neurons that receive excitatory vs. inhibitory (or mixed) signals varied, depending on the route that the DCN signals followed: the CB-VTA pathway was more likely to communicate excitatory signals to neurons in NAc_Core_ than in NAc_Med_, whereas the CB- thalamic pathways did not show a clear preference for either NAc subregion.

Our neuroanatomical studies identified three disynaptic pathways for CB-NAc communication through the VTA and intralaminar thalamus (CM, PF) nodes. Disynaptic pathways that utilize fast classical neurotransmitters (e.g., glutamate) would be expected to support rapid signal propagation and result in short-latency responses, whereas neuromodulatory (e.g., dopaminergic) pathways would be expected to operate at longer time scales (Chen et al., 2014; Gao et al., 2018; Assous et al., 2019; Liu et al., 2021; Yang et al., 2021; Sippy and Tritsch, 2023). The shortest-latency responses we recorded upon photostimulation of cerebellar axons in each of the intermediary nodes occurred within 10 ms, a finding that is consistent with the monosynaptic connections known to exist between DCN and VTA, and DCN and thalamus (Carta et al., 2019; Baek et al., 2022; Jung et al., 2022), and which agrees with our newly identified structural connectivity. Particularly for responses elicited by photostimulation in VTA, DCN axons are known to synapse onto both dopaminergic and non-dopaminergic VTA neurons (Beier et al., 2019; Carta et al., 2019; Baek et al., 2022), including vesicular glutamate transporter 2-positive (vGlut2+) VTA neurons, which project to NAc (Beier et al., 2015) and which could, at least partly, mediate cerebellar signaling. Longer-latency excitatory responses dominate CB-NAc communication through the intralaminar thalamus and could arise from polysynaptic circuitry, in agreement with the diffuse patterning of intralaminar projections (Jones and Leavitt, 1974; Van der Werf et al., 2002). Subthreshold synaptic signaling that requires activity-dependent short-term facilitation in order to propagate to NAc neurons could also account for longer latencies. Given that optogenetic stimulation of CB axons with a single pulse could elicit NAc responses, short-term plasticity does not appear to be strictly required for successful CB-NAc communication. However, it could account for the excitatory NAc responses evoked through PF, which displayed more gradual buildup than excitatory responses through the other nodes.

Inhibitory responses are also a key component of CB-NAc connectivity and may be important for global gain control (Buzsáki et al., 2007). Inhibitory responses (and/or inhibitory components of mixed responses) were seen upon stimulation in DCN, or in any of the nodes, and occurred with longer latencies compared to excitatory responses. They also appeared to depend on photostimulation intensity, with higher prevalence at lower photostimulation intensities. Multiple cellular mechanisms could underlie inhibitory responses and could involve dopamine release as well as non-dopaminergic signaling through the VTA (Brown et al., 2012; Beier et al., 2019; Carta et al., 2019; Baek et al., 2022) and/or CB-thalamic (CM, PF) projections to NAc, which are known to contact not only medium spiny neurons but also cholinergic inhibitory interneurons (Yamanaka et al., 2018). Polysynaptic pathways could also be involved, although their recruitment seems at odds with the inverse relationship between inhibitory response prevalence and photostimulation intensity. The nature of synaptic signaling and plasticity in CB-NAc circuits is an exciting route for further investigation.

### Technical considerations

Our recordings were performed in ketamine-anesthetized mice. Although ketamine at anesthetic doses does not alter the number of spontaneously active striatal neurons (Kelland et al., 1991), it affects the pattern of spontaneous activity, promoting step-like membrane potential fluctuations between “up-” and “down-states” (Mahon, 2001). Such bursting behavior would contribute to “noisy” baseline activity; however, we would not expect it to distort our analyses because we have defined neural responses as changes in spiking rate from the preceding baseline. Moreover, by using electrical and/or optogenetic, rather than sensory, stimuli to probe connectivity, we have overcome any concerns stemming from the known inhibitory effect of ketamine on sensory-evoked activity in the cerebellum (Bengtsson and Jörntell, 2007).

In addition to anesthesia, the stimulation modality (electrical, optogenetic) warrants consideration. Electrical stimulation could potentially result in antidromic activation of DCN inputs, which could lead to recruitment of alternate polysynaptic pathways to NAc. This concern was mitigated by our optogenetic experiments, in which ChR2-expessing DCN axons were specifically activated in the nodes. Moreover, although electrical current propagates widely in the brain, the use of bipolar electrodes allowed us to restrict the spread of DCN stimulation to ∼100 µm, as shown by the substantial drop in the probability to activate the same NAc neuron with DCN stimuli delivered 100 µm apart. Lastly, optogenetic stimulation of DCN axons in a node could induce back-propagating action potentials that could, in turn, elicit NAc responses through axonal collaterals. However, to the best of our knowledge, there is no evidence for DCN collaterals that form synapses onto NAc-projecting neurons in both VTA and intralaminar thalamic nuclei. Even if such collaterals existed, their activation would result in unreliable responses with relatively long latencies, due to synaptic delays and the failure-prone asymmetric nature of action potential conduction velocity (Grill et al., 2008; Mateus et al., 2021).

### Novelty of anatomical connections

Anatomical and functional connections between DCN and VTA, and DCN and intralaminar nucleus nodes, have been described previously (Snider and Maiti, 1976; Hendry et al., 1979; Phillipson, 1979; Watabe-Uchida et al., 2012; Gornati et al., 2018; Carta et al., 2019; Jung et al., 2022). VTA and thalamic projections to NAc are also well established (Beckstead et al., 1979; Su and Bentivoglio, 1990; Bentivoglio et al., 1991; Bassareo and Di Chiara, 1999; Garris et al., 1999; Groenewegen et al., 1999; Van der Werf et al., 2002; Ikemoto, 2007; Taylor et al., 2014). It might therefore be unsurprising that the DCN recruit these nodes to communicate with NAc. However, the VTA and thalamus are exquisitely complex areas with multiple output streams. For example, the VTA projects not only to NAc but also to hippocampus, hypothalamus, lateral habenula, entorhinal cortex, etc. (Oades and Halliday, 1987; Stamatakis et al., 2013; Beier et al., 2015). None of these downstream target regions appears to be disynaptically connected to DCN through the VTA, which points to specificity in circuit wiring. Our study is the first one, to our knowledge, to map DCN-NAc circuits through VTA and thalamus. A recent study provided anatomical evidence for a medial DCN-NAc circuit (Fujita et al., 2020). Here we extend these observations to lateral DCN and also provide evidence for the centromedial and parafascicular nuclei as parts of the thalamic node.

### Ideas and Speculations

It is tempting to speculate on network-wide implications of the different CB-NAc pathways. Fast excitatory signals would be well poised to support the rapid communication of information critical to the control of motivated behavior, such as prediction or prediction-error signals, which are well established in the cerebellum (Wagner et al., 2017; Heffley and Hull, 2019; Kostadinov et al., 2019; Larry et al., 2019). The fact that DCN signals can reach NAc_Core_ and NAc_Med_ through distinct anatomical (and possibly cellular) pathways leaves room for distinct multiplexing with signals originating elsewhere.

Cerebellar signals that arrive to NAc_Core_ through VTA could be behaviorally relevant for reward learning through signaling of value, which has been correlated with dopamine release in NAc_Core_, but not in NAc_Med_ (Mohebi et al., 2019). Alternatively (or additionally), the CB-VTA- NAc_Core_ pathway could be rewarding in itself, as stimulation of dopaminergic projections from VTA to NAc_Core_ are sufficient to establish self-stimulation (Han et al., 2017) and previous studies have shown that the CB can signal reward (Wagner et al., 2017; Carta et al., 2019; Medina, 2019). The CB-VTA-NAc_Core_ pathway could also be involved in social cognition, given that NAc_Core_ activation promotes social dominance in mice and that cerebellar Purkinje neurons are known to modulate social aggression in mice (Jackman et al., 2020). Finally, the CB-VTA- NAc_Core_ pathway could also be important for signaling salience, regardless of valence, to promote learned responses to behaviorally relevant stimuli (Kutlu et al., 2021). The short response latencies of the CB-VTA-NAc_Core_ pathway could be key for temporally precise control of behavioral states including attentional control (Flores-Dourojeanni et al., 2021), and could gate incoming signals to NAc, including signals from PF (Akaike et al., 1981; Hara et al., 1989).

The observed bias of CB-PF neurons to preferentially influence NAc_Med_ over NAc_Core_ could support a role in behavioral flexibility. Indeed, both NAc_Med_ and NAc_Core_ neurons encode rewarding and aversive stimuli via changes in neural firing rate, however, only neurons in NAc_Med_ adaptively shift population response type to track the motivational value of a stimulus (Loriaux et al., 2011), suggesting that NAc_Med_ can flexibly modulate encoding based on relative stimulus valence and/or acquired salience (Aquili et al., 2014; West and Carelli, 2016). Given that the PF is known to play a role in behavioral flexibility (Brown et al., 2010; Kato et al., 2021), the CB-PF-NAc_Med_ circuit may communicate teaching signals for updating past learning and altering behavior adaptively.

Future investigations into the cellular basis of DCN-NAc communication as well as potential circuit-specificity of behavioral contributions are clearly in order. Here, we have broken new ground by providing the first evidence of rapid functional connectivity between CB and NAc, through the midbrain and intralaminar thalamus.

## Materials and Methods

### In vivo electrophysiology

### Mice

C57Bl/6J mice of both sexes were used in accordance with the National Institutes of Health guidelines and the policies of the University of California Davis Committee for Animal Care. Acute in vivo recordings were performed at postnatal days P43-P91. All animals were maintained on a light/dark cycle (light 7 am - 7 pm), and experiments were performed during the light cycle.

### Surgery

Mice of both sexes (P35 - P49; N = 11) were used for optogenetic experiments. Animals were anesthetized with isoflurane (4% - 5% induction; 1.5% maintenance) and secured to a stereotactic frame (Kopf Insruments, Tujunga, CA). After exposing the top of the skull, the head was leveled and small craniotomies were drilled over the DCN. Channelrhodopsin-expressing adeno-associated virus (AAV2-hSyn-hChR2(H134R)-EYFP, UNC Vector Core, Chapel Hill, NC; 100 nl, 5.6x10^12^ gc/ml, 1:2 dilution) was injected into the three DCN using a Micro4 controller and UltraMicroPump 3 (VWR, Radnor, PA). Glass needles were made from 1-mm outer diameter glass pipettes (Wiretrol II, Drummond Scientific Company, Broomall, PA) pulled to a fine tip (20 - 50 µm tip diameter, 3 - 4 mm tip length) using a pipette puller (P-97, Sutter Instrument, Novato, CA). Injection needles were left in place for 5- 10 min following injections to minimize diffusion. The following coordinates were used for targeting pipettes to each nucleus (relative to lambda): medial n.: -2.55 AP, +/-0.8 ML, -2.15 DV; interposed n.: -2.45 AP, +/-1.7 ML, -2.15 DV; lateral n.: -1.82 AP, +/- 2.37 ML, -2.17 DV. Following surgery and analgesia administration (0.1 mg/kg buprenorphine, 5 mg/kg meloxicam), mice were allowed to recover on a warm heating pad before being transferred back to the vivarium. Animals were given 4-5 weeks expression time before being used in in vivo recording experiments.

For electrophysiological recordings, anesthesia was initially induced by brief inhalation of 4-5% isoflurane followed by an intraperitoneal injection of anesthetic cocktail (100 mg/kg ketamine; 10 mg/kg xylazine; 1 mg/kg acepromazine) and was maintained with periodic injections of the anesthetic cocktail (20-50 mg/kg), as needed. After confirmation of anesthesia depth using a toe pinch response test, mice were placed in a stereotactic frame (Stoelting Co., Wood Dale, IL) on a heating pad. Breathing rate and toe pinch responses were monitored to ensure maintenance of anesthesia. For electrical stimulation experiments (N = 42 mice of both sexes), a small craniotomy and durectomy were performed over the DCN (lateral/interposed n.: relative to lambda, in mm: -2.1 to -2.6 AP; +/-2.1 to +/-2.3 ML). For optogenetic experiments, optic fibers were implanted over the CM (relative to bregma: -1.3 AP; -.85 ML; - 3.55 DV, 10° angle), PF (relative to bregma: -1.87 AP; -.95 ML; -2.95 DV, 12° angle), or VTA (relative to bregma: -3.1 AP; -0.5 ML; -4.05 DV) ipsilaterally to recording site and secured with temporary resin acrylic (Keystone Industries, Gibbstown, NJ). For recordings, craniotomy and durectomy were made over NAc, targeting the medial shell and/or core (relative to bregma: +1.7 to +1.54 AP; +/- 0.4 to +/-1.15 ML).

### Electrical/Optical Stimulation

For electrical stimulation, a custom stereotrode (∼200 μm distance between tips) was lowered 1.95 - 2.45 mm below the brain surface (in DV axis) through the cerebellar craniotomy to reach the DCN. Ten trials of bipolar constant-current electrical microstimulation were delivered at each location at a 15-s inter-trial interval. Each stimulation trial consisted of a 200- Hz burst of five 0.5-ms monophasic square-waveform pulses at 100 μA. In a subset of experiments (N = 15 mice), the stimulation intensity was varied between 30, 100, and 300 μA in interleaved blocks of 5 trials per intensity. When stimulation was repeated at different DCN sites in the same experiment, an effort was made to move the stereotrode in a direction perpendicular to the axis of the two poles, so that an at least partly distinct pool of cerebellar neurons could be stimulated. For optical stimulation, 20-50 trials of ten 5-ms light pulses (473- nm DPSS laser, Opto Engine LLC, Midvale, UT) were delivered at 20 Hz (inter-trial interval: 30 s or 1 min). Optical stimulation was primarily delivered at 10 mW intensity at fiber tip, but lower intensities were also tested (1 and 5 mW).

### Acquisition of Electrophysiological Data

Recordings were performed with a 12-channel (platinum; 0.2 - 2 MΩ impedance) axial multi-electrode array (FHC Inc., Bowdoin, ME; 150 μm distance between channels), dipped in fluorescein dextran (3000 MW) and lowered into the NAc at a depth of 3.75 - 5.25 mm below the brain surface (DV axis). Electrode signals were fed to a digital headstage (RHD 2132, Intan Technologies, Los Angeles, CA) for amplification (x20), filtering (0.7 - 7500 Hz) and digitization (30 kS/s with 16-bit resolution), before transfer to an open-source acquisition system (OpenEphys) for display and storage.

### Histology for Verification of Electrode/Optic Fiber Placement

Positioning of electrodes and fibers was initially guided by atlas-based stereotactic coordinates (Paxinos & Franklin) and, upon completion of experiments, histologically verified through electrolytic lesions (for location of DCN microstimulation electrodes; single 10-s cathodal pulse of 300 µA), fiber track inspection (for optic fiber) and fluorescence imaging (for recording array; NAc fluorescein dye track). Coordinates for subsequent animals of the same litter were further adjusted accordingly. At the end of experiments, electrodes were retracted and animals were perfused with 4% (w/v) paraformaldehyde (PFA) in 0.1 M phosphate buffer (PB). Brains were dissected and post-fixed in 4% PFA, sliced in phosphate buffered saline (PBS) and inspected under a fluorescence stereoscope (Olympus, Tokyo, Japan; SZX2). NAc slices containing the dye track from the recording array were identified under 488 nm light; CB slices with electrolytically lesioned tissue, and thalamic and/or VTA slices with optic fiber tracts, were identified under brightfield illumination. Slices of interest were subsequently stained with DAPI (1:20,000, Thermo Fisher Scientific Inc., Waltham, MA), mounted on slides, and imaged using a VS120 Olympus slide scanner. Images were manually registered to the Paxinos and Franklin Mouse Brain Atlas and the location of the recording electrode tip and optic fiber were mapped. Only channels along the recording array that were determined to be within NAc_Core_ and/or NAc_Med_ were included in analysis. Experiments with misplaced optic fibers, viral expression outside the DCN region, or cerebellar electrolytic lesions localized outside the lateral and/or interposed DCN, were excluded from analysis.

### Quantification and Statistical Analysis Electrophysiological data processing and quantification

Custom-written MATLAB scripts (Mathworks, Natick, MA) were used for data processing and analysis. Filtering of spikes was achieved through a 4^th^-order Butterworth high-pass filter (cutoff at 300 Hz). The filtered signal was thresholded at 3-3.5 standard deviations below the average voltage. Spikes were subsequently sorted in putative single-unit clusters with Plexon Offline Sorter (version 3.3.5, Plexon Inc., Dallas, TX) using principal component analysis. All well-isolated neurons were included in the analyses. Firing rates were computed from spike counts in consecutive 10-ms time bins (peri-stimulus time histogram; PSTH). Spikes with time differences less than 7 ms from stimulus artifacts or 1 ms from the previous spike were discarded.

Quantification of stimulation-induced modulation of NAc spiking activity followed recent studies of cerebellar neurons (Chen et al., 2014; Washburn et al., 2022) and our own evaluation of the dataset. Specifically, we averaged the PSTH across all trials of the same intensity for any given pair of recording and stimulation sites and normalized to the average pre-stimulus baseline (z-score). We defined an excitatory response as an increase in firing rate that exceeded 3 standard deviations (σ) above the baseline within the first 500 ms after the stimulus onset. An inhibitory response was defined as a decrease in firing rate of more than 1 σ below the baseline that persisted for a minimum of three time bins (i.e., 30 ms) and started within the first 500 ms after stimulus onset. The 500-ms time window captured more than 90% of responses (not shown). For single-unit responses that included both excitatory and inhibitory components (mixed responses), we quantified each component individually for amplitude and latency. For illustration purposes only, all firing rates were convolved with a Gaussian kernel (σ = 16 ms). Response onset latency was defined as the first instance at which firing rate changed from baseline as described above, using 1-ms time bins. For analyses of multi-unit activity, see our BioRxiv preprint (D’Ambra et al., 2020).

### Statistics

Statistical analyses were performed in Graphpad Prism 9.3.0 and Matlab. Comparisons of the relative frequency distribution of response types were performed with Contingency Table analyses (Chi-square tests). Comparisons of response latencies and firing rates were performed with t-tests and ANOVA. For permutation tests, we randomly shuffled the data between groups 1,000 times and estimated the probability to find a difference greater than or equal to the observed difference by chance alone (Oden and Wedel, 1975; Antzoulatos and Miller, 2011). To correct for multiple comparisons, we followed the Benjamini-Hochberg method (Benjamini and Hochberg, 1995) with False Discovery Rate (FDR) set at 10%.

### Surgery for anatomical studies

For anatomical tracing, juvenile mice of both sexes were injected with AAVs and tracers using the procedure described above for in vivo optophysiology. For co-localization experiments: AAV9-CAG-GFP (2 x 10^12^ viral particles/ml, UNC viral core) was injected in DCN (from bregma, in mm: medial n.: -2.55 AP, ± 0.75 ML, -2.1 DV, 50 nl; interposed n. : -2.5 AP, ± 1.55 ML, -2.1 DV, 50 nl; lateral n.: -2.2 AP, ± 2.3 ML, -2.12 DV, 50 nl; and -1.8, ± 2.35 ML, -2.12 DV, 50 nl). Cholera toxin subunit B (ctb)- 640 or -568 (5 mg/ml, Biotium) was injected in NAc medial shell and core (from bregma, in mm: 1.8 AP, ± 0.8 ML, -4.2 DV, 200 nl). For anterograde transsynaptic tracing: AAV1-hSyn-Cre-WPRE-hGH (10^13^ gc/ml, Addgene; 1:10 dilution) was injected in DCN (coordinates as above). AAV5-CAG-FLEX-tdTomato (7.8 x 10^12^ viral particles/ml, UNC viral core; 1:2 dilution) or AAV5-pCAG-FLEX-EGFP-WPRE (1.1 x 10^13^ gc/ml, Addgene; 1:2 dilution) was injected in VTA (from bregma, in mm: 2.8 AP, ± 0.35 ML, - 4.2 DV, 100 nl; and -2.85 AP, ± 0.6 ML, -4.2 DV, 100 nl) or thalamus (from bregma, in mm: - 0.85 AP, ± 0.3 ML, -3.3 DV, 300 nl; and -1.2 AP, ± 0.5 ML, -3.5 DV, 300 nl; 1:5 dilution). Following surgery, mice remained in the colony for 2-3 weeks to allow for recovery and retrograde labeling/virus expression prior to euthanasia and tissue collection/processing.

### Histology and fluorescence microscopy for anatomical studies

Two to three weeks following tracer/virus injections, mice were anesthetized with the anesthetic cocktail and perfused transcardially with 4% (w/v) PFA in PB. Brains were post-fixed in 4% PFA for 6 h and transferred to 30% sucrose in PBS for overnight incubation at 4°C. Brains were coronally sectioned (60 μm) on a sliding microtome, stained with DAPI, mounted to slides, coverslipped with Mowiol-based antifade solution and imaged. VTA sections were immunostained for tyrosine hydroxylase (TH) prior to mounting, as follows: slices were first incubated with blocking solution [10% normal goat serum (NGS) in PBS supplemented with 0.3% Triton-X100; PBST] for 1 h at room temperature, then with mouse anti-TH (clone LNC11, Millipore Sigma, Burlington, MA; 1:1000) in blocking solution with 2% NGS overnight at 4°C. Sections were washed with PBST (3 x 20 min) and incubated for 1 h at room temperature with goat anti-mouse Alexa fluor 488 secondary antibody (1:1000), washed with PBS (3 x 20 min), mounted, coverslipped and imaged. Epifluorescence image mosaics were acquired on an Olympus VS120 slide scanner with a 10x air objective. High magnification confocal images were taken sequentially with different laser lines and a 63x oil-immersion objective on a Zeiss LSM800 microscope with Airyscan. Image brightness/contrast was adjusted using ImageJ (NIH) for display purposes. Data from a total of N = 13 mice with successful injections, without spill to neighboring regions, are presented.

## Acknowledgments

We thank Amber Arif and Alicia Dye for help with tissue sectioning; Drs. Brian Wiltgen, Karen Zito, Marty Usrey and Brian Mulloney of UC Davis for access to equipment; Dr. Ditterich for comments on a previous version of the manuscript; and Dr. Chaudhuri for helpful discussions. Author contribution: AD, EA and DF designed the research; AD, SJJ and KV performed the research; AD, KV and EA analyzed the data; SG assisted with tissue slicing and registration; SJJ prepared the anatomy figures; AD, EA and DF wrote the paper with feedback from all authors. This work was supported by a Whitehall fellowship, BRFSG-2017-02, R21MH114178, NSF1754831 and R01MH128744 to DF; a NARSAD 2018 Young Investigator Grant to EA; AD was supported by NIH T32 GM007377 and a UC Davis Dean’s Distinguished Graduate Fellowship; KV was supported by NIMH T32 MH112507 and F31MH131405 by NIMH.

